# Analysis of an open source, closed-loop, realtime system for hippocampal sharp-wave ripple disruption

**DOI:** 10.1101/298661

**Authors:** Shayok Dutta, Etienne Ackermann, Caleb Kemere

## Abstract

Transient neural activity pervades hippocampal electrophysiological activity. During more quiescent states, brief ≈100 ms periods comprising large ≈150–250 Hz oscillations known as sharp-wave ripples (SWR) which co-occur with ensemble bursts of spiking activity, are regularly found in local field potentials recorded from area CA1. SWRs and their concomitant neural activity are thought to be important for memory consolidation, recall, and memory-guided decision making. Temporally-selective manipulations of hippocampal neural activity upon online hippocampal SWR detection have been used as causal evidence of the importance of SWR for mnemonic process as evinced by behavioral and/or physiological changes. However, though this approach is becoming more wide spread, the performance trade-offs involved in building a SWR detection and disruption system have not been explored, limiting the design and interpretation of SWR detection experiments. We present an open source, plug-and-play, online ripple detection system with a detailed performance characterization. Our system has been constructed to interface with an open source software platform, Trodes, and two hardware acquisition platforms, Open Ephys and SpikeGadgets. We show that our *in vivo* results — approximately 80% detection latencies falling in between ≈20–66 ms with ≈2 ms closed-loop latencies while maintaining <10 false detections per minute — are dependent upon both algorithmic trade-offs and acquisition hardware. We discuss strategies to improve detection accuracy and potential limitations of online ripple disruptions. By characterizing this system in detail, we present a template for analyzing other closed-loop neural detection and perturbation systems. Thus, we anticipate our modular, open source, realtime system will facilitate a wide range of carefully-designed causal closed-loop neuroscience experiments.

## 1 Introduction

Historically, precisely-targeted lesion studies have been the gold standard to establish causal relationships between neural regions and cognitive functions, for example studies showing hippocampal involvement in recognition and Pavlovian contextual fear memories [1, 2]. Modern tools such as opto- and chemo-genetics enable these perturbations to have reversible effects and to be targeted to specific genetically-specified target cells. However, many neural circuits are now understood to exhibit distinct patterns of activity at different times which are hypothesized to serve specific functions. Causal manipulations to illuminate these functions require the ability to interact with the brain — detect and perturb neural activity — in closed loop. Often times, these patterns are transient events of neural activity comprised of sub-second “ensemble events”. In order to modulate information contained in these patterns, they must be detected not only with high accuracy but also low latency.

Closed-loop systems for detecting and perturbing temporally-distinct patterns of neural activity obligatorily combine both algorithmic (i.e., software) and computational hardware components. There is often a naive assumption that accuracy can be maximized and latency minimized via computational or hardware improvements. However, in many regimes, it is possible that algorithmic trade-offs may be the primary factors constraining detection latency and accuracy. Without a system-level analysis, however, these trade-offs may remain poorly-understood and investigators lose the opportunity to balance performance against potential side-effects of neuromodulation such as seizures, tissue damage, or the induction long-term plasticity. There are two primary challenges to system-level analyses. First, closed-source data acquisition platforms limit the ability to understand or explore the full data processing pipeline. Second, parameter explorations during on-going *in vivo* experiments are limited by the finite nature of neural implants and animal cooperation.

We present an open-source system we developed to detect sharp-wave ripples (SWRs) — brief periods of elevated ≈150-200 Hz oscillation in the local field potential (LFP) which are prominent in area CA1 of the hippocampus — and trigger perturbations of ongoing activity (e.g., silencing activity via optogenetic stimulation to hippocampal area CA3 or electrical stimulation of the ventral hippocampal commissural fibers). Such systems have recently been used to investigate the causal significance of neural activity during SWRs in learning, consolidation, and working memory [3–7]. In order to investigate our realtime system, we explored algorithmic parameter spaces by employing a two-level strategy. First, we developed a simulation model of the system allowing us to rapidly and efficiently prototype algorithms and evaluate performance. Second, we built an apparatus to feed pre-recorded neural activity back into our data acquisition system and ran pseudo-experiments with varied parameters. We applied this strategy to both a synthetic “gold-standard” dataset and to data recorded *in vivo* from hippocampal area CA1. After identifying key performance metrics, we show how varying parameters affects system performance using an initial single-channel detection algorithm. We demonstrate how these performance trade-offs depend on both algorithmic choices and system hardware. Finally, we show how enhancing the algorithm to use multiple detection channels can improve system performance at a cost of only minimally increasing processing load and latency.

## 2 Methods

### 2.1 Animal Use

For the *in vivo* portion of our study, one male Long Evans rat (Charles River Laboratories) was implanted with a micro-drive array with independently adjustable tetrodes (implant coordinates −3.66 mm AP and 2.4 mm ML relative to bregma). Tetrodes were lowered into hippocampal CA1 over one week. Tetrode locations were determined by characteristic LFP waveforms attributed to the target area in conjunction with estimated tetrode depth. A skull screw was used as a ground while a tetrode left within the corpus callosum picking up minimal neural activity was used as a digital reference and subtracted away from all tetrodes. We let the rat explore an open field with objects and hidden rewards prior to all recording sessions within a sleep box. All experiments were approved by the Rice University Institutional Animal Care and Use Committee’s guidelines and adhered to National Institute of Health guidelines.

### 2.2 System Framework

Our experimental system architecture, depicted in Figure 1, is a closed-loop system beginning with data acquisition and ending with a digital pulse triggered upon realtime SWR detection. This pulse may be used to trigger a ripple disruption mechanism (e.g., optogenetic fiber or biphasic current stimulator). Data is passed through a headstage to an Open Ephys or SpikeGadgets data acquisition unit, which interfaces with a software suite, Trodes, for which we developed a SWR detection module. Upon detection of a SWR event, the module triggers a stimulation with a digital pulse via an Ethernet-connected BeagleBone Black microcontroller.

**Figure 1:**
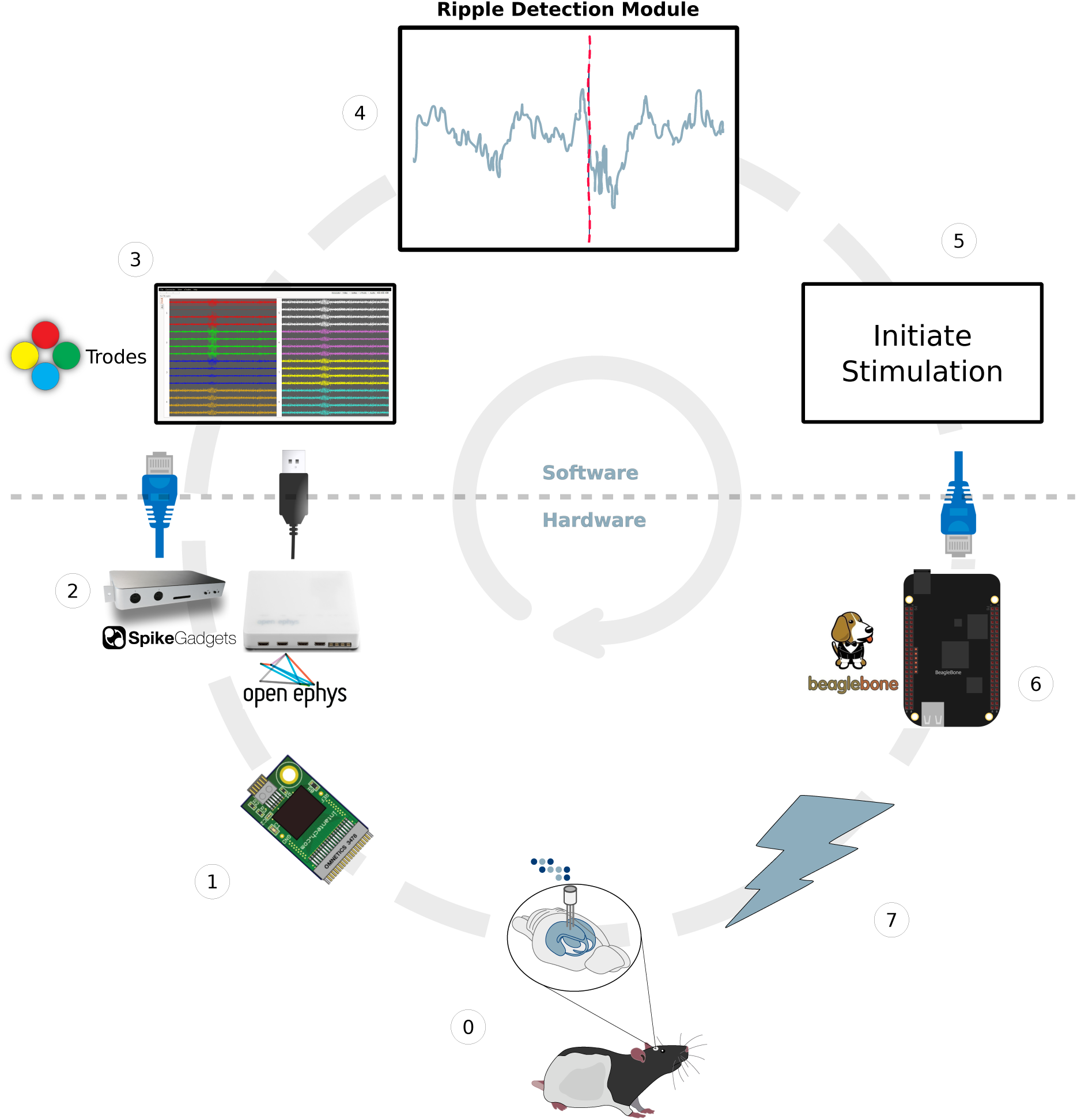
Closed-loop system architecture depicting LFP data acquisition, data transfer (blue Ethernet cables and black USB cable), data processing, and stimulation. Process begins with data acquisition and ends with selective stimulation based on sharp-wave ripple detection. Figure adapted from Open Ephys.

#### 2.2.1 Data Acquisition

LFP data is collected through tetrodes which connect to a headstage that digitizes the raw analog LFP signal via an onboard analog-to-digital converter. This digitized signal is then transmitted to a computer for display and/or further signal processing. As depicted in Figure 1, step 2, we used two different data acquisition systems, the Open Ephys Acquisition Board and SpikeGadgets Main Control Unit, to acquire the digitized signal at 30 kHz from Intan RHD2000 series headstages. These systems differ in the protocol used to transmit data to the computer — the SpikeGadgets Main Control Unit employs gigabit Ethernet, whereas the Open Ephys Acquisition Board uses USB2.0. Note that the USB buffer size, is configurable and affects round-trip latency. We found the lowest round-trip latency from 30 kHz data collection to feedback from the BeagleBone Black to be when the number of USB blocks to read was set to three. The two hardware systems evaluated are developed by two organizations, Open Ephys (open-ephys.org) and SpikeGadgets (spikegadgets.com), that promote custom interactions with neural data through a variety of open-source software and tools enabling novel neuroscience experiments.

#### 2.2.2 Data Processing and Stimulation Trigger

Trodes — a modular, open source, cross platform software suite available from SpikeGadgets (spikegadgets.com/ software/trodes.html) for neural data acquisition and interaction and experimental control — was used for data collection and signal processing. The software is written in QT/C++ enabling various realtime applications. As depicted on the left side of Figure 2, Trodes is a multi-threaded process with threads interfacing with the data acquisition unit and sending data for realtime display and logging (Figure 1 step 3). Incoming data from both the headstage and available auxiliary channels are sampled at 30 kHz with both acquisition units in this study and handled by the appropriate Data Acquisition threads. The Stream Processor thread then sends filtered data as requested by connected modules. This modular framework facilitates custom realtime applications as various forms of data (wide-band electrical signals, spike data, animal position, etc.) can be sent to the module processes using Trodes.

The SWR (or ripple) detection module, shown in Figure 1 step 4 and on the right side of Figure 2, receives digitized signals (discussed in the subsection above) of specified channels via UDP communication protocol from the Trodes Stream Processor thread. The digitized signals are sent to the module after a 400 Hz low-pass filter (LFP band) and decimation from 30 kHz to 3 kHz. The module process receives the data via the Trodes Data Interface thread and passes it along to a central thread, Stimulation Handler, which spawns off separate threads per channel utilized for ripple detection (algorithm discussed in Detection Algorithms subsection). Once enough channels report ripple detections within a certain time requirement (15 ms for our experiments) to the Stimulation Handler, a separate thread initiates stimulation by communicating with a BeagleBone Black microcontroller over Ethernet. The microcontroller outputs a 3.3 Volt digital pulse for a user specified amount of time (100 microseconds in this study) to trigger a disruption mechanism for closed-loop intervention. However, as this paper evaluates the efficacy of the detection system, for our analysis we monitor the timestamps at which the disruption pulse is triggered by feeding it into the auxiliary digital inputs of the acquisition units as opposed to stimulation hardware.

**Figure 2:**
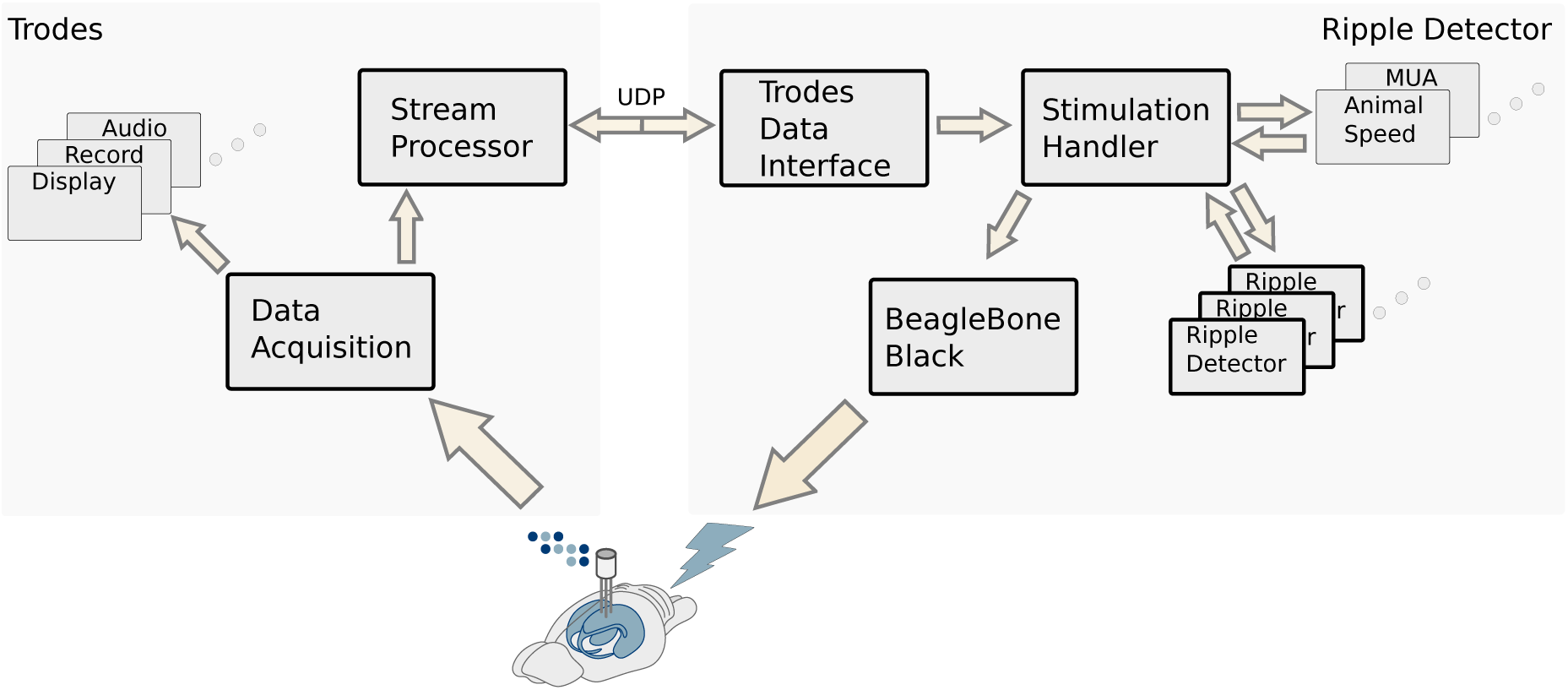
Trodes and Ripple Detector processes (highlighted on the left and right, respectively) with their various threads. Data transfer and processing threads of importance in this study are bolded. Trodes process collects, logs, and transfers data as requested by connected modules. The Ripple Detector module processes incoming data and triggers a potential stimulation pulse upon event detection.

### 2.3 Detection Algorithms

#### 2.3.1 Canonical Ripple Detection

Post-recording, ripple events were defined on tetrodes that displayed characteristics of the CA1 area of the hippocampus. Specifically, the recorded LFP in one of the channels of the selected tetrode (same one subject to online detection for our realtime analysis) first had a digital reference subtracted away. This signal was then LFP band filtered with a 400 Hz low-pass infinite impulse response (IIR) filter (from Trodes). Afterwards the signal was decimated and ripple band filtered (150–250 Hz) with a 25 tap finite impulse response (FIR) filter. Ripple band filtering was done using a forward and a time-reversed path, resulting in a net zero group delay (time shift from filtering). The instantaneous power of the ripple band filtered signal was then calculated via a Hilbert Transform and further smoothened with a Gaussian kernel with a 4 ms standard deviation. Ripple events were detected as times when z-score of the smoothened power signal signal exceeded a threshold of 3 z-units for at least 15 ms. The canonical ripple epochs were defined as the time points from which the processed signal returned down to the mean before and after threshold crossings [8, 9]. In cases when multiple electrodes (typically channels on different tetrodes) are available for ripple detection, a different canonical definition is required. Ripples were initially defined using as above for each electrode. A canonical multichannel ripple was defined as one which is simultaneously detected on each electrode (two in our analysis). The multichannel ripple epoch is defined as the union of the detected single-channel ripple epochs, i.e., the start of the earliest ripple detected and to end with bound of last ripple detected. As such, we obtain a conservative ripple detection latency estimate while covering the entire span of the time the LFP is in a high ripple band power state. We reanalyzed our data with the canonical ripples being defined on different channels and tetrodes with a 300 tap bandpass FIR filter allowing 1% “ripple” in the passband with −30 dB suppression in the stopband but our results and subsequent conclusions remained consistent.

#### 2.3.2 Realtime Ripple Detection

Like our canonical ripple detections, the realtime or online detection algorithm is comprised of single channel and multichannel modalities. The single channel case performs realtime reference subtraction from the LFP (filtered in the same way as the offline case) and decimation from 30 kHz to 3 kHz as in the canonical detection case. However, the difference between online and canonical detections begins from ripple band filtering. The decimated signal is filtered to the ripple band with a 30 tap FIR filter. In order to perform a realtime instantaneous power estimation and smoothing, the realtime algorithm computes the absolute value of the ripple band filtered signal and further filters it by a 33 tap 50 Hz low-pass FIR (instead of a Hilbert transform followed by Gaussian kernel smoothing). These filters cause an intrinsic sample delay from the offline case (≈10.167 ms in our case). It is worth noting that the number of filter taps as well as filter types were determined by analyzing algorithmic delay and detection accuracy based on metrics described in the Data Analysis subsection. To normalize detection thresholds in the realtime case, the mean and standard deviation of the smoothed envelope are estimated over a 20 minute training period. In two ≈90 minute sleep box recording sessions, when we sampled 20 minute time intervals at random (N=1000), the resulting mean and standard deviation were within 5% of the values for the entire sessions. This length of time and subsequent error in parameter estimation likely depends on the behavioral and/or sleep state of the animal — in our experimental recordings, animals were contained in a sleep box. realtime detections are then triggered when the envelope crosses a threshold defined as α standard deviations above the mean (*threshold* = α *∗ σ* + µ) or α z-units. Following a detection, there is a 200 ms lockout period where we ignore any further threshold crossings (i.e., to avoid stimulation artifacts). Additionally, we impose a hard limit on the number of detections per second (set to three during the experiments in this work). For multichannel detection, the single channel algorithm runs on separate threads per electrode. A specified number of channels upon which ripples are being detected must all be above threshold within 15 ms of each other in order to trigger a detection event. In the multichannel case, the lockout is imposed following the multichannel detection event. In order to precisely characterize our system, we additionally implemented an offline simulator which carried out the same algorithmic steps in a sample-accurate way. This allowed us to differentiate between algorithmic and hardware, communication, and/or operating system delays.

### 2.4 Realtime Testing

#### 2.4.1 Synthetic

Prior to *in vivo* testing, we developed a synthetic “gold-standard” dataset to validate efficacy of our algorithm and to determine baseline latency quantifications. To replicate CA1 neural noise dynamics within the ripple frequency bands (150–250 Hz), we generated a white noise process and then filtered it to the ripple band. This signal was then adjusted to have the same standard deviation as the ripple band filtered CA1 LFP recording. Ripple events were then injected into the synthetic signal. Synthetic ripples were generated by multiplying a 200 Hz sinuosoid with a Gaussian envelope. The envelope was chosen to have ≈100 ms standard deviation length (average length of ripple events from our recordings), with a peak value equal to the average maximum peak of ripples found within our recording. This process generated a synthetic ripple event which was then added to the previously generated synthetic neural noise giving us a dataset of synthetic “gold-standard” ripples. The final dataset used in our testing was 15 minutes long with 500 injected ripples.

The dataset was then converted to a .wav audio file and played through the realtime system via custom PCB shorting all channels of the headstage to an auxiliary cable input. This framework was used for testing the system with both Open Ephys and SpikeGadgets data acquisition hardware. However, different headstages were used for testing these two systems. A 32 channel headstage was used with the Open Ephys hardware while a 128 channel headstage was used with the SpikeGadgets hardware. Digital pulses corresponding to ripple detections, parameter values of the detection algorithm, and wide-band synthetic data across all channels, were logged during each data collection session. It is worth noting that different headstages have inescapable intrinsic noise fluctuations during the recordings. Data was reanalyzed across different channels and using different headstages at different points in time validating our results and subsequent conclusions.

#### 2.4.2 In vivo

For *in vivo* testing of our system, a rat was placed inside a sleep box and then tethered to the SpikeGadgets Main Control Unit via a SpikeGadgets HH128 headstage. Data was recorded through our detection module for ≈60-90 minutes in three separate sessions in order to test two different modalities of our algorithm (discussed in Detection Algorithms subsection) and for synthetic data generation. As in the synthetic case, ripple detection pulses, parameter values of the detection algorithm, and wide-band electrophysiological data across all channels, were logged during each data collection session. Lastly, it is worth noting that prior to the recording sessions, the rat explored an open field with novel objects and hidden rewards in order to increase the prevalence of SWRs [10–14].

### 2.5 Data Analysis

We characterized system and algorithmic performance via a total of five different metrics: true positive rate (TPR), false positive percentage, false positive/stimulation rate (FPR), detection latency, and relative detection latency. All metrics, other than false positive percentage, were evaluated on both synthetic and *in vivo* data. Metrics for *in vivo* analyses were generated by first binning the an ≈80 minute dataset into 15 ms chunks followed by random sampling with replacement of these chunks until a total of ≈80 minutes had been sampled. The metrics generated were then averaged and this bootstrapping process was repeated 1000 times. Metrics for the synthetic data analysis, however, were calculated over the entire dataset without any bootstrapping or averaging. As we controlled the duration of the synthetic dataset as well as the number of canonical ripples, a false positive percentage metric as opposed to rate was used to characterize performance. Metrics used were calculated as follows:

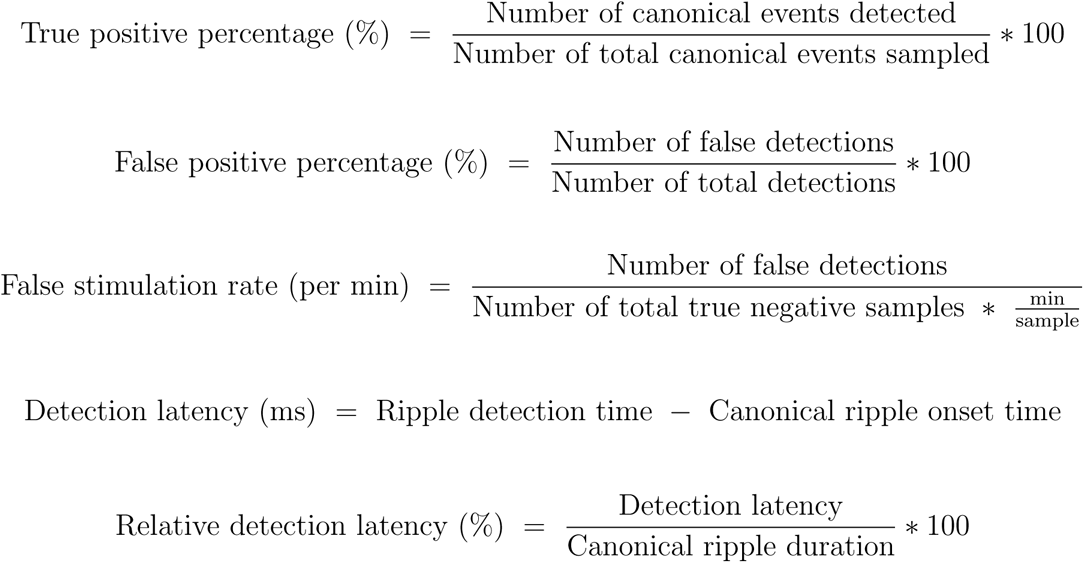

In our evaluations of the realtime system, as well as simulated detections, we ignore canonical ripples that had start times within 200 ms of the previous putative ripple since 200 ms is the post disruption lockout period that we use (see Detection Algorithms subsection). All analysis was done with custom Python scripts, a modified version of the Nelpy package, and custom C++11 programs. Both analysis code and figure data are publicly available at https://www.github.com/shayokdutta/RippleDetectionAnalysis.

## 3 Results

For closed-loop experiments which seek to disrupt the neural activity that occurs during or in response to SWR, there are three figures of merit: we want to maximize true positive rate, minimize false positive rate and minimize latency. Why are these important? In a closed-loop perturbation experiment, failure to detect a substantial fraction of events will increase the likelihood that no effect of perturbation will be detected. A high rate of false positive-induced perturbations will contaminate experimental results and can potentially cause harm (i.e., seizures). Finally, for most experiments detection latency is just as important as detection accuracy – perturbations that do not perturb the desired patterns while they are going on will not have the intended effect. Using synthetic and actual data, we demonstrate below that these figures of merit are linked and describe their trade-offs for single and multichannel ripple detection.

The SWR detection system we developed and analyzed runs as a module in Trodes (Figure 1, 2). Two features in particular enabled our system analysis: First, as our neural recording system digitizes neural recordings with appropriate anti-aliasing filters, the recorded signal could be converted back to the original analog form modulo a constant scale factor. This enabled us to repeatedly replay identical data into the system to test parameter settings. Second, access to the complete source code — not only of our SWR detection module but also the underlying data acquisition system enabled us to simulate each step of the data processing pipeline. Our module constructs a realtime representation of ripple-band (150–250 Hz) power through a simple FIR filter followed by taking the magnitude of the filtered signal and further smoothing through a low-pass FIR filter (see Methods and Figure 2). Using statistics of the ripple-power signal learned during an initial training period to normalize, it then detects SWR events as excursions in estimated power over a defined threshold. Note that each of these processing steps deterministically delays the neural signal in ways which depend on the parameters of the algorithm. This work investigates the trade-offs associated with choosing these parameters, particularly the value of the threshold.

We built a detailed simulation of our ripple detection module which carried out the same sample-by-sample computations and produced identical values at each step as the online system. We used synthetic and recorded hippocampal test data to evaluate performance. Using these test data, our algorithm simulation enabled us to quantify how variability in the ripple signal deterministically transformed into variability in detection performance. In parallel, converting the test data back to analog form, we fed them into the actual system and re-digitized the signals including the timestamp of the generated ripple disruption pulse. By comparing the simulated and actual timestamps of ripple detection we could quantify the statistics of the non-deterministic contributors to performance which result from use of a general purpose computer for signal processing. This process enabled us to broadly explore the parameter spaces and extrapolate realtime system performance before actual closed-loop testing *in vivo* (Figure 6).

### 3.1 Latency and Accuracy trade-offs with Synthetic Data

Variability is one of the greatest challenges in optimizing the detection of particular signals in the brain. In particular, SWR events vary in amplitude, duration, precise frequency content, etc., and while individual large exemplars are easy to identify, even trained investigators have difficulty in determining the boundary between smaller SWR events and spontaneous high frequency oscillations. Thus, in order to confirm the validity of our realtime algorithm, we created a gold-standard synthetic ripple dataset. This synthetic data was formed by adding 500 ripple events to 15 minutes of background noise. While real ripple events vary in peak amplitude and duration, we used a constant peak amplitude and duration for synthetic ripple events to eliminate the effect of this variation on system performance. This allowed us to analyzed performance as a function of peak amplitude.

Our *in vivo* recording sessions showed a high percentage of ripples with peaks ranging from approximately 5 to 10 z-units in the normalized ripple band power signal. As such, our “gold-standard” ripple library consisted of distinguishable ripples varying in peak size around this range relative to a constant level of background activity (see Methods). Our module detected ripples both in simulation and via playback to our hardware systems. The detection threshold in the algorithm is specified in terms of z-units — standard deviations above the mean — of the estimated envelope of the underlying synthetic ripple band noise. In the case of large ripples (e.g., Figures 3A and 3B), a detection threshold exists which results in detection of all synthetic ripples and for which incorrect detections (false positives due to background noise) never occur. In the example shown, ripples have a peak power of 10 z-units, and the threshold is set to 5 z-units. When actual events lie closer in power to the background activity, however, it is clear that it will be impossible to select a threshold for which no true events are missed and no false positives are accidentally detected. We quantified this effect by sweeping detection threshold values in datasets generated to contain synthetic ripples with different peak powers.

**Figure 3:**
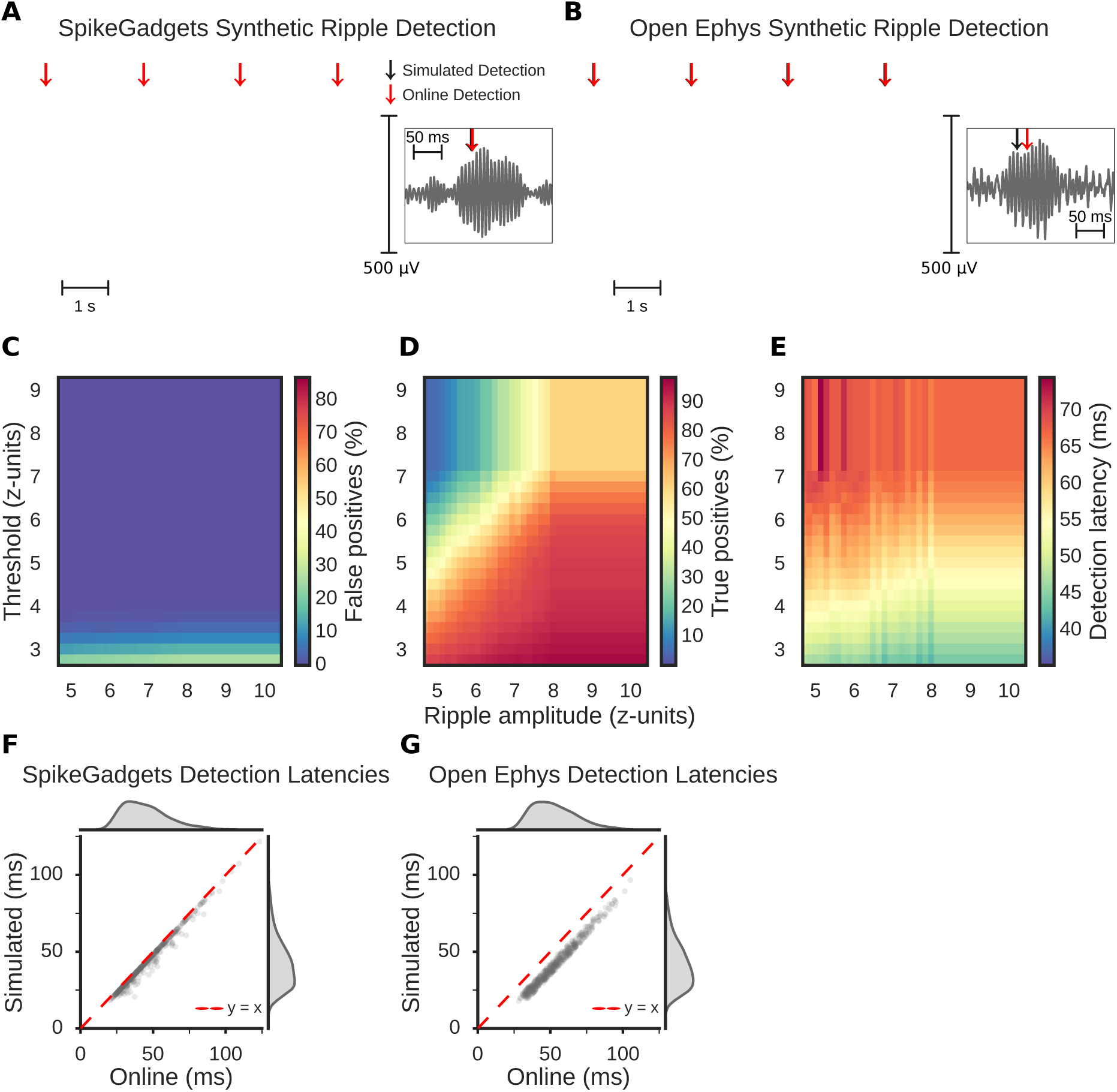
**(A, B)** Example detections using SpikeGadgets and Open Ephys, respectively. Canonical ripples are highlighted and both simulated and online detection times marked (black and red arrows, respectively). **(C–E)** Performance characterizations with synthetic ripples of varying amplitudes and simulated online detections with varying thresholds. **(C)** False positive percentages reveal threshold at which synthetic ripples exceed baseline noise. **(D)** True positives taper down with increasing threshold across ripple amplitudes. **(E)** Detection latency heatmap shows general increase in latency with increasing threshold across ripple amplitudes. **(F, G)** Simulated vs online detection latency scatter plots and distributions for synthetic ripples sent through SpikeGadgets and Open Ephys hardware, respectively.**(F)** Using SpikeGadgets, average latencies were 42.79 ms and 45.44 ms for simulated and online, respectively, with 80% closed-loop latencies lying within 1.35 and 2.6 ms. **(G)** Using Open Ephys, average detection latencies were 41.65 ms and 53.32 ms for simulated and online, respectively, with 80% of closed-loop latencies lying within 7.5 and 13.8 ms.

Across our synthetic datasets, the power of the background noise was held constant. Thus, the false positive percentage depended only on the chosen threshold (Figure 3C). In contrast, detection requires that events be larger than the threshold, yielding a surface where for a given threshold detection accuracy is dependent on ripple amplitude or events will be missed (Figure 3D). Both physiological ripples and our synthetic events have a non-zero rise time. Hence, as the threshold rises, the latency — the delay from when the ripple started (exceeded the mean of the background power) and when it is detected — increases (Figure 3E). Our synthetic ripples were generated using a Gaussian envelope which was scaled by peak power. The result is that larger ripples rise in power faster, and thus are detected with lower latency for a given threshold. As a result, we see that the detection threshold jointly affects latency and accuracy (true and false positive), and thus the optimal setting will depend on the details of a particular experiment.

### 3.2 Realtime Detections with Synthetic Data

Our simulation revealed that choosing a particular value of threshold would yield ripple detections with specific latency and accuracy trade-offs. However, these trade-offs assume that signal processing occurs instantaneously. While some custom-hardware ripple interruption implementations may approach this limit, the need for flexible design has led many investigators to use general purpose computers to carry out computation. In this scenario, communication with hardware and other operating system functions leads to further delay between a ripple occurring and detection yielding a closed-loop response.

In order to quantify these non-algorithmic effects on performance, we proceeded to feed in synthetic data to our realtime system. After estimating a threshold of 5 z-units online by feeding in 2 minutes of background noise, we ran the 15 minute synthetic ripples dataset with 500 injected ripples. As expected, realtime detections of individual synthetic ripples were identical to our simulations, but with increased latency (deviation from the unity-slope line, Figure 3F and 3G). We found that the additional, realtime latency, was different for our two data acquisition systems (80% of the closed-loop latencies were within 1.35–2.6 ms and 7.5–13.8 ms for SpikeGadgets and OpenEphys, respectively). Given that the system pipeline is identical following data acquisition, this suggests three things. First, in the case of the computations required for ripple detection, time spent computing the algorithm is at most a few milliseconds. Second, the specific interface protocol used for data acquisition, i.e., USB or Ethernet (Figure 1 step number 2), can significantly impact the closed-loop latency. Lastly, this isolates that the majority of overall ripple detection latency lies in the algorithmic implementation —filter-delay and the time it takes for the processed signal to reach threshold — rather than in hardware or computational efficiency. The synthetic analysis as a whole highlighted key performance characteristics of our detection algorithm, validated the efficacy of our realtime system, and identified a speedup with the Ethernet-based SpikeGadgets Main Control Unit — used exclusively for the *in vivo* portion of our study.

### 3.3 In Vivo Single Channel Detection Analysis

Our investigation of system performance using synthetic data demonstrated that for different amplitude ripples, there would be a trade-off between accuracy and latency. Physiological ripples vary in intensity and duration and *in vivo* recordings can also contain bursts of high-frequency noise induced by static discharge, animal behaviors such as grooming, whisking, or chewing. How does this variability affect the performance of a realtime ripple detection system? We began answering this question by applying our system simulation to recorded *in vivo* data. We recorded neural activity from 10 tetrodes implanted in area CA1 of the hippocampus during an ≈80 minute session in which a rat rested in a small box. Unlike synthetic data, physiological ripples cannot be identified perfectly. There is no universally agreed definition; we applied a canonical definition of ripples as periods in which the power in the ripple band (150–250 Hz) was elevated above the mean by 3 standard deviations for at least 15 ms. We defined the start and end of each ripple epoch as the moment when the ripple-band power first exceeded and finally returned to the mean (see Methods). As discussed previously, the detection threshold in our realtime detection algorithm is also specified in terms of standard deviations of the envelope of the ripple signal. For our *in vivo* parameter explorations, we used mean and standard deviation parameters calculated over the entire 80 minute session.

As with synthetic data, while increasing the threshold parameter, we found that true positives and false positives decreased with increasing threshold (Figures 4A–4B). For real data, as we do not know the number of ground truth events, we calculate a false positive or false stimulation rate to give us an informative characterization of how long we are disrupting information propagation from the hippocampus over a period of time. As expected, latency increased with increasing threshold (Figures 4C–4D). Unlike in our synthetic dataset, not all real ripples are of constant length. As such, the absolute detection latencies only partially characterize the performance of our detection algorithm. In order to fully interpret these values, we incorporated a relative latency metric — the percent of the ripple event that has transpired prior to detection (see Methods).

**Figure 4:**
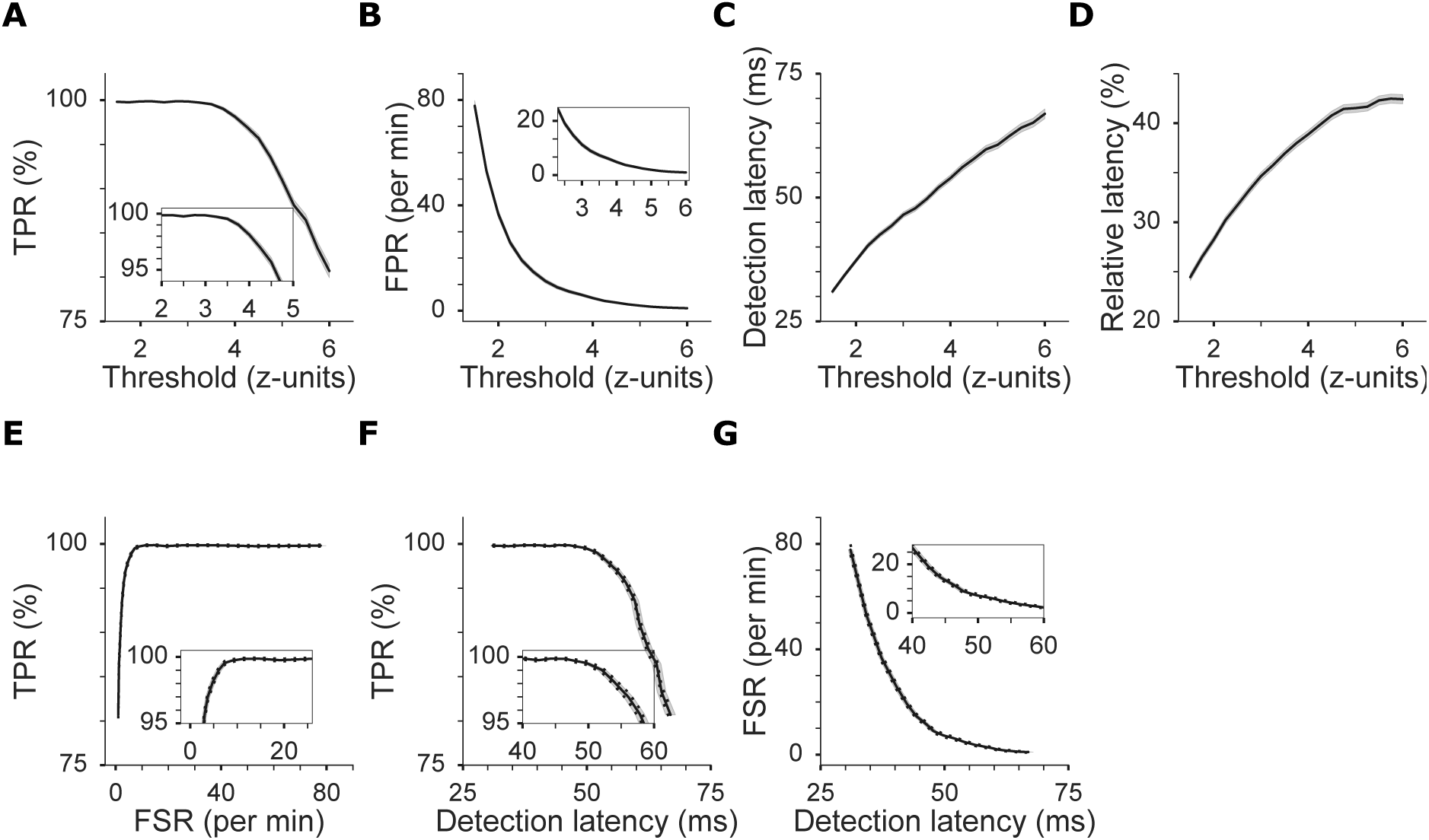
Single channel *in vivo* ripple detection metrics. Mean and two standard errors of data points (solid and shaded regions, respectively) are shown in **(A–D) —** generated using a Monte Carlo sampling approach (see Methods). **(A)** True rate (TPR) decreases as threshold increases. **(B)** False positive rate (FPR) decreases with increasing threshold. **(C)** Detection latency increases with increasing threshold. **(D)** The relative detection latency, the fraction of each event that has transpired prior to detection, also increases with increasing threshold. Panels **(E–G)** show mean and two standard errors of data points. The dashed lines represent standard errors on the y-axis while the shaded region represents standard errors in the x-axis. **(E)** Receiver operating characteristic (ROC) curve shows true positive and false positive percentage relationships. **(F, G)** False stimulations decrease with as detection latency increases but true positives also decrease with increasing detection latency.

We additionally visualized the pairwise trade-off curves relating true and false positive rates and detection latency Figures 4E–4G). Unlike with our synthetic data, the false stimulation metric is non-zero for all threshold values (Figure 4B) and there is a larger range for which true and false positive rates trade off as shown in the receiver operating characteristic (ROC) curve in Figure 4E.

Depending on the specifics of experimental design, one can imagine that extraneous perturbations (i.e., false positives) or missed events (i.e., false negatives) might be more detrimental. In many cases, however, investigators would want to pick a threshold value associated with the “elbow” of the ROC (inset in Figure 4E) which is associated with minimizing false positives while maximizing true positives. Our results indicate that given between 5–10 false positives detections per minute, greater than ≈97% of the canonical ripple events can be detected. The threshold parameter space associated with this region lies in between ≈3.5 and 4.25 z-units (Figures 4A and 4B).

Prior to selecting a parameter associated with the “elbow” of the ROC curve for realtime testing, the corresponding latencies must also be considered. The latency values associated with the 5 to 10 false detections per minute and the greater than ≈97% true positive detections range fall in between ≈45–55 ms (Figures 4F and 4G). These latency values further isolate the desired threshold range to fall within a similar range as the true positive and false positive trade-offs (Figures 4A - 4C). We see in Figure 4D that given these suggested performance metrics, choosing threshold values up to 4 or even 4.25 can lead to both sensitive and specific interaction with up to ≈60% of the ripple event.

### 3.4 In Vivo Two Channel Detection Analysis

When we examined false positive detections in real data, we noted that a common signature was a brief period of minimally-elevated ripple band power that did not meet the duration criteria. The fact that physiological ripples propagate across CA1 has previously inspired closed-loop ripple detection systems which require a ripple to be detected on multiple electrodes [4, 15]. We implemented and evaluated an algorithm that required that the realtime power estimate be above threshold on two selected channels within 15 ms in order to trigger detection, referred to as the multichannel or two channel algorithm. The canonical ripple definition was updated to use ripples with temporal overlaps across multiple channels (see Methods). For comparison, we analyzed both individual electrodes using the single channel algorithm, but with the new multichannel canonical definition (Figure 5).

**Figure 5:**
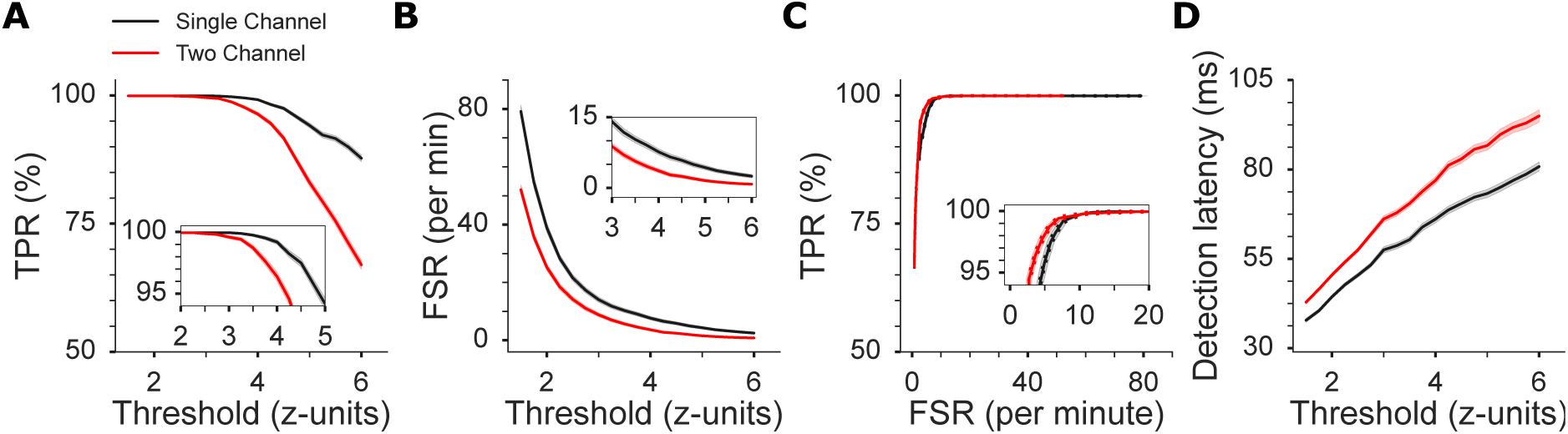
Single channel and two channel in vivo ripple detection metrics. Note that two channel detections provide more accurate detection results than single channel at lower thresholds. Threshold/metric definitions and all plots display same metrics and same trends as in Figure 4 but with a two channel comparison and ground truth ripples based on two channel detections (see Methods).

**Figure 6:**
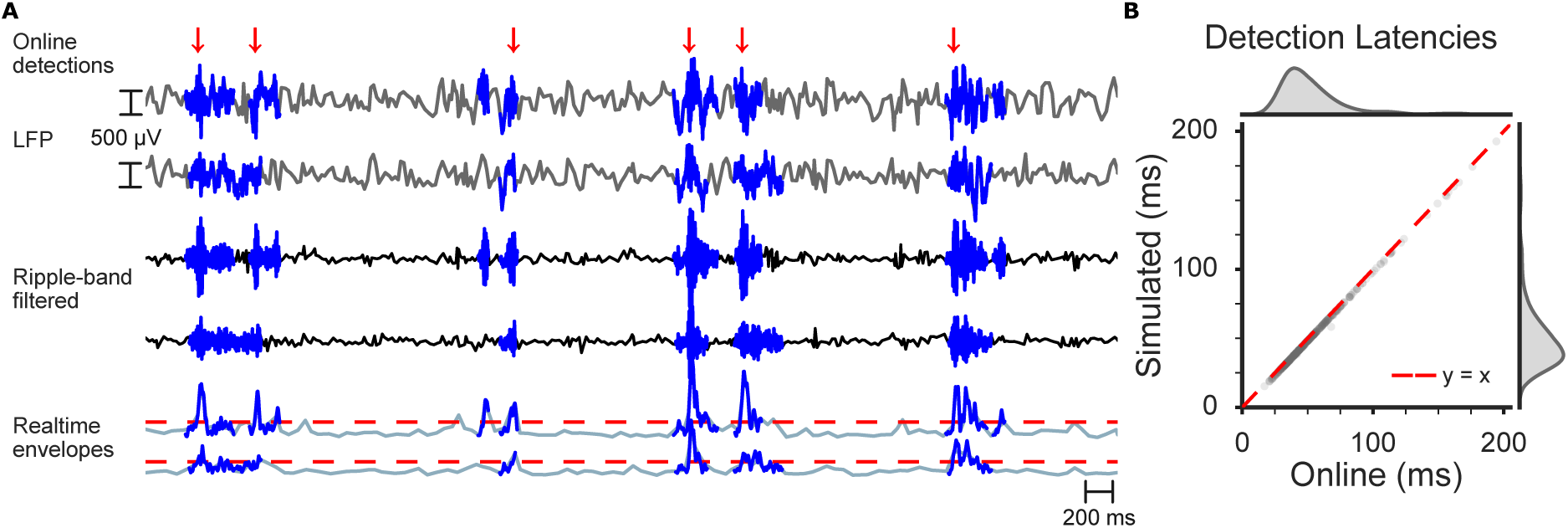
**(A)** Example snippet of two channel in vivo ripple detections. Blue highlights across all channels mark canonical ripple epochs. Dark blue colored traces in each channel represent individual ripple detections per channel done offline. The red horizontal traces represent realtime threshold of 3 z-units on both channels used for the detection. **(B)** Scatter plot of detection latencies in between simulated detections (threshold crossing times) and online detections. Average simulated and online detection latencies are 53.322 ms and 54.621 ms, respectively. Average and median latencies between threshold crossing time and online detection time are 1.924 ms and 2 ms, respectively.

The more stringent requirement that a ripple be present on multiple channels results in a more rapid decrease in the fraction of true positives detected compared with single channel detection (in which the true multichannel ripple need only be detected on one channel, Figure 5A). As expected, this decreased sensitivity comes with the benefit of increased selectivity (Figure 5B). However, the two channel algorithm can be as sensitive as the single channel algorithm at lower thresholds with lower FPRs. For example, a threshold of 3.25 z-units with the two channel voting algorithm results in a higher TPR than single channel detections with a threshold of 4.25 z-units while having comparable average FPRs (insets, Figures 5A and 5B). This is further evinced in the ROC curve which combines true and false positive metrics (Figure 5C).

Intuitively, we expect the two channel algorithm to have higher detection latencies at similar thresholds as it waits for a second channel to detect a ripple prior to triggering a detection pulse. We indeed find this (Figures 5D). Thus, the increased accuracy of multichannel detection comes at the cost of increased latency. This, and the fact that ripples (unlike theta oscillations) are not coherent across the whole hippocampus, suggest that care should be taken in the development of multichannel detection systems [15, 16]. As the numbers of channels is increased beyond two, our module features a custom-voting algorithm, as opposed to unanimous voting, which requires a user specified voting requirement of ripple detections prior to a stimulation pulse output which may increase sensitivity to investigator preferences [4, 17].

After our offline and simulated analysis, we proceeded to implement the multichannel algorithm in our realtime system. With a rat inside a sleep box for approximately 60 minutes, we ran an *in vivo* two channel voting and detection session. We then set the detection threshold to 3 z-units after estimating mean and standard deviation parameters for 20 minutes (see Methods). According to the two channel analysis, at this threshold we should be able to detect greater than 97% of all ripple events and detection latencies in between 50–55 ms on average. Figure 6A shows a snippet of the recorded LFP and realtime processed signals. Note the temporal lag in the processed signals (see Methods). Two of the single channel ripple detections, traced in dark blue, are not considered part of the canonical ripple epochs, vertically highlighted across all channels. The online detections selectively ignore these events and only detects the canonical ones as they cross threshold across both the channels used for the detection within 15 ms (shown in the bottom realtime envelope traces). Lastly, as we did while passing our synthetic data through the realtime setup, we examined algorithmic performance in both simulated and online cases. Figure 6B shows the distributions of both online and synthetic detections of the same ripple events during this recording session. We are able to confirm that both our average online and simulated detection latencies, 54.621 ms and 53.322 ms, respectively, fall within the range of our synthetic analysis of this recording session with the given parameters. Additionally, on average the closed-loop latency is ≈2 ms indicating that adding another channel is not causing significant computational delay compared to single channel. This was further confirmed by running the online detection algorithm with eight channels voting for ripple detections on our synthetic dataset. We once again found a similar closed-loop latency of approximately 2 ms on average (not shown).

## 4 Discussion

Within the last decade, electrical and optogenetic ripple disruption studies have established a fundamental role for SWRs in the processes of learning and memory [4, 18–20]. However, performance characterizations of realtime SWR detection systems have not been discussed. Here, we have introduced and evaluated an open source, closed-loop, realtime system for SWR detection and potential intervention. Our study began with generating a “gold-standard” synthetic ripple dataset, revealing subsets of z-scores of ripple band power associated with underlying neural activity and SWRs. We then proceeded to test our algorithm and system performance with the synthetic dataset prior to *in vivo* testing which motivated further algorithmic improvements and isolated particular pitfalls of our detection algorithm.

For preliminary evaluations of algorithmic performance, we generated a synthetic ripple dataset as ground truth neural data is an underlying challenge in neuroscience. Our online estimated power-threshold crossing algorithm showed that the algorithm was valid in detecting synthetic ripples that were injected into our dataset. However, as mentioned in the Methods section, the fabrication of the synthetic dataset involved modeling the underlying neural noise within the ripple band. Comparing z-scores of the Hilbert Transform of this signal to the CA1 LFP revealed that the real data encapsulated the modeled noise. We found that the CA1 z-scores that had a longer tail than the modeled noise (z-scores of ≈5 and above) were associated with ripple events. As a result, when we injected synthetic ripples into our dataset that had a larger power within the ripple band than the surrounding noise, it was expected that our detection algorithm would be able to selectively detect those events and would be valid criteria for preliminary testing of the realtime system (Figure 3E).

Testing the realtime setup with a synthetic dataset also identified a closed-loop latency speedup by using SpikeGadgets hardware over Open Ephys for data acquisition (Figure 3F–3G). Our results revealed an added latency from simulated to online detections due to computational time and hardware latency. We discovered the latter to be the primary source of the closed-loop latency by demonstrating that adding another thread for detections in our two channel detections (Figure 6B) from our single channel detections (Figure 3F) did not increase the simulated to online detection time latency (the gap from the *y* = *x* line). Additionally, we stated the 8 ms speedup by using a SpikeGadgets acquisition unit over the Open Ephys acquisition unit was due to the use of an Ethernet over USB communication protocol. Specifically, the USB protocol involves putting data into packets and then sending those packets with a variable number of blocks of data making up the packets. We varied the number of USB blocks, which increased and decreased the latency, to further confirm that the use of this protocol was the reason for the speedup offered by Open Ephys. The results we presented were found by using the number of USB blocks that provided the lowest latency. It is worth noting that the Open Ephys project is developing a platform for data transfer via PCI express which will enable sub-millisecond closed-loop latency [22]. Lastly, to further confirm that computational processing load was not the primary factor in the added closed-loop latency, we tested our realtime system with 8 simultaneous channels detecting synthetic ripples and confirmed that our closed-loop latencies using both systems provided results consistent with single channel detections. This investigation and demonstration further confirmed our claims about the closed-loop latency being a factor of hardware-software interfacing rather than computational time. In general, we believe that using either data acquisition unit will enable closed-loop ripple disruption experiments as added hardware and computation latencies are <10% of calculated relative detection latency of ripples.

Moving from synthetic ripple detection to *in vivo* introduced increased false detections due to ripple events sharing ripple band power z-scores with underlying neural activity and noise events from rodent grooming or whisking. Real ripples, unlike our model, do not always reach a value upon which they can be distinctly isolated from other events. This makes the resultant false detections from a single channel threshold crossing algorithm inevitable. In an attempt to lower false detection metrics, we evaluated a multichannel detection algorithm founded on our understanding that ripple events should permeate multiple tetrodes within CA1. Multichannel detections resulted in a lower total number of ripple detections and higher selective detections based on our accuracy metrics than relying upon a single channel (Figure 5). However, unfortunately, both ripple and noise events were coherent across tetrodes. Additionally, events of ripple band power that exceeded threshold without meeting the temporal requirements of our canonical ripples were also prevalent across tetrodes. Using two channels for detections did lower the number of detections due to these neural dynamics but it did not fully address the issue. The remaining challenge involves false detections due to noise events.

As noise events saturate all tetrodes, a proposed solution to handle false detections due to rodent grooming or static passed through to the data acquisition unit would be to utilize a false detection tetrode located in a brain region picking up minimal activity. This false detection channel would perform ripple detections as the previously discussed detection channels but the digital output pulse from our realtime system would be suppressed if this channel reports a detection a few milliseconds before or after (buffer window) the other channel(s) report ripples [23]. Preliminary analysis of this paradigm reveals that false detections due to noise events associated with the reference channel as well as due to rodent grooming or other sources can indeed be suppressed without drastically impacting the true positive rate. However, the trade-off of such an algorithm comes with the extra buffering required after ripple detection. Investigators will have to determine what sort of sensitivity is appropriate given the latency cost of the buffer window.

The other source of false detections, neural dynamics within the LFP associated with ripple band power exceeding thresholds, remains an open question to be solved. Several examples can be found in all of our datasets where ripple band power exceeds thresholds set and triggers our realtime detections but do not meet the 15 ms above threshold temporal requirement of canonical ripples. From the perspective of neural dynamics within the hippocampus, the general purpose of detecting ripples remains to disrupt periods of high multiunit activity co-occurring with sequential reactivation or replay. As such, without proper identification of replay during population bursts, presenting an inherent challenge itself, it may not be truly accurate to call these events false detections. While we have not discussed it here, our module includes the ability to interface with a video tracking system and limit perturbation to periods in which animal speed is below a threshold value.

Nevertheless, based on our definitions and data analysis metrics, it is possible to algorithmically lower the number of false detections while maintaining high true positive percentages. One of the differences between our online algorithm compared to the offline canonical ripple definition is the temporal threshold requirement (i.e., above threshold for at least 15 ms). The temporal requirement was dropped in the online algorithms due to added detection latency for time spent after threshold crossing in addition to the closed-loop latency associated with any realtime system. As previously discussed, we found our relative detection latencies for both modalities of our algorithm to vary between 35%–45% (Figures 4D) at thresholds of interest. Additionally, we found that ≈95% of ripples observed within our study last from ≈60–150 ms with a mean at ≈100 ms. This indicates that every ms of above-threshold temporal requirement added would increase relative detection latency by ≈1% of the ripple’s duration. Further investigations and analysis of thresholds while adding temporal requirements in realtime may provide a choice in parameters that result in detections that better align with putative ripple events. Overall, the proposed realtime temporal requirement may raise detection latencies into the 60%−70% range, limiting us to interacting the latter 30%−40% of the event.

Adding in a temporal requirement for online detections and increasing relative detection latencies leads into a question of the meaning manipulating the latter percentage of a ripple event. The hippocampus is known to communicate with the neocortex for memory consolidation resulting in memories being more dependent on cortical regions than the hippocampus [21, 24, 25]. As such, it may be possible that the percentage of the ripple event that has occurred prior to intervention is consistently being propagated through and we are limited to affecting a certain amount of cognition. This may potentially explain why goal-related ripple disruption experiments, requiring full sequential reactivation of particular sequences, have been shown to impair performance while disruptions during non-sequential tasks (e.g., environmental recognition) tend to have more limited effects [4, 6, 18–20]. Ultimately, further studies and investigations into neural content disrupted upon realtime SWR detection compared to the subset of activity that propagates through to downstream cortical regions are required in order to understand the effects of disruption and purposes served by SWRs.

## 5 Conclusion

Altogether, our work investigated the parameter space of ripple detections and identified the trade-offs between true and false positive percentages along with detection latency. For any ripple-power based detection system, It appears that a threshold in the range of 3 to 4.5 standard-deviations above the mean will optimize performance for most experiments with a rodent inside a sleep box. The choice of single or multichannel detection will depend on the number of available electrodes and the extent of the perceived importance of detecting coherent events across large fractions of the hippocampus. We anticipate that the low latency, realtime, closed-loop, SWR detection system we implemented in the open-source neural data acquisition software suite, Trodes may be widely useful, particularly in conjunction with low-cost, open-source data acquisition units from Open Ephys in enabling the dissemination of this experimental technique. With our system investigators have the ability to further fine-tune the threshold to detect desired events observed online while using our analysis as a guide. Finally, as the number of closed-loop perturbation experiments increases, we hope that our simulation and validation approach will inspire careful exploration of parameter and performance metric spaces.

## Acknowledgements

This work was supported by National Science Foundation (CBET-1351692 and IOS-1550994), the Human Frontiers Science Program (RGY0088) and the Ken Kennedy Institute for Information Technology. We are grateful for the assistance of past and present Realtime Neural Engineering lab members during data acquisition. Additionally, we would like to thank Chun-Ting Wu, Matthijs van der Meer and members of the van der Meer lab for beta testing our realtime SWR detection system.

